# Marmosets as model systems for the study of Alzheimer’s disease and related dementias: substantiation of physiological Tau 3R and 4R isoform expression and phosphorylation

**DOI:** 10.1101/2024.04.26.590453

**Authors:** Hasi Huhe, Sarah M. Shapley, Duc Duong, Fang Wu, Seung-Kwon Ha, Sang-Ho Choi, Julia Kofler, Yongshan Mou, Thais Rafael Guimaraes, Amantha Thathiah, Lauren K.H. Schaeffer, Gregory W. Carter, Nicholas T. Seyfried, Afonso C. Silva, Stacey J. Sukoff Rizzo

## Abstract

**INTRODUCTION:** Marmosets have been shown to spontaneously develop pathological hallmarks of Alzheimer’s disease (AD) during advanced age, including amyloid-beta plaques, positioning them as a model system to overcome the rodent-to-human translational gap for AD. However, Tau expression in the marmoset brain has been understudied.

**METHODS:** To comprehensively investigate Tau isoform expression in marmosets, brain tissue from eight unrelated marmosets across various ages was evaluated and compared to human postmortem AD tissue. Microtubule-associated protein tau (*MAPT*) mRNA expression and splicing were confirmed by RT-PCR. Tau isoforms in the marmoset brain were examined by western blot, mass spectrometry, immunofluorescence, and immunohistochemical staining. Synaptic Tau expression was analyzed from crude synaptosome extractions.

**RESULTS:** 3R and 4R Tau isoforms are expressed in marmoset brains at both transcript and protein levels across ages. Results from western blot analysis were confirmed by mass spectrometry, which revealed that Tau peptides in marmoset corresponded to the 3R and 4R peptides in the human AD brain. 3R Tau was primarily enriched in neonate brains, and 4R enhanced in adult and aged brains. Tau was widely distributed in neurons with localization in the soma and synaptic regions. Phosphorylation residues were observed on Thr-181, Thr-217, and Thr-231, Ser202/Thr205, Ser396/Ser404. Paired helical filament (PHF)-like aggregates were also detected in aged marmosets.

**DISCUSSION:** Our results confirm the expression of both 3R and 4R Tau isoforms and important phosphorylation residues in the marmoset brain. These data emphasize the significance of marmosets with natural expression of AD-related hallmarks as important translational models for the study of AD.

## INTRODUCTION

Non-human primates provide critical insight into primate-specific mechanisms that are the etiologies of human diseases. Given the translational limitations of rodent models, the common marmoset (*Callithrix jacchus*) has emerged as an important model system for studying diseases of aging, including Alzheimer’s disease (AD) [1–5]. The marmoset brain has conserved neuroanatomical and neurocircuitry to that of humans [6, 7], as well as age-related cognitive decline and neuropathological features that align with postmortem tissues of human AD patients[1, 8, 9] One of the major pathological hallmarks of AD, amyloid-beta (Aβ) deposition in the brain has also been reported frequently in marmosets as early as 7 years of age, which is equivalent to a 56-year-old human [8]. Despite the well-reported characterization of Aβ in the aging marmoset brain, only a few studies have comprehensively characterized Tau in the marmoset brain [9–12].

Tau is a microtubule-associated protein encoded by the *MAPT* gene, predominantly expressed in the brain, and plays essential roles in neuronal function. During development, alternative splicing of *MAPT* results in 6 isoforms of Tau transcripts in humans, which include 0N3R, 1N3R, 2N3R, 0N4R, 1N4R, and 2N4R. In humans, the 0N3R Tau isoform is exclusively expressed in fetal and neonatal brains, while alternative splicing of exon 10 results in the expression of both 3R and 4R isoforms in the adult brain, which are maintained throughout the lifespan with an almost equimolar ratio under physiological conditions [13–16]. Under pathological conditions, including as a consequence of neurodegeneration, alterations of exon 10 splicing occur, resulting in an altered 3R/4R Tau ratio, which is associated with distinct Tauopathies, including in AD [17–20]. In humans, physiological Tau is not only localized in axons but also distributed in dendritic spines at synaptic terminals where it plays essential roles in synaptic plasticity[21–25], including long-term depression (LTD) formation, which is necessary for cognitive function[26, 27]. In the AD brain, synaptic Tau accumulates and forms oligomeric Tau [21], which has been shown to impair synaptic function [28–30] via trans-synaptic propagation of Tau pathology [31, 32]. The formation of pathological Tau aggregates is tightly associated with post-translational modifications of Tau, especially hyperphosphorylation [33]. It is well established that Tau hyperphosphorylation plays an essential role in facilitating pathological Tau aggregation[34] and that phosphorylation residues change during AD progression. For example, Thr-181, Thr-217, and Thr-231 hyperphosphorylation has been reported in the early preclinical stages of AD [35], while AT8 (Ser202/Thr205) and PHF1(Ser396/Ser404) hyperphosphorylation are reported in later stages of AD [36, 37].

While several studies have characterized Tau extensively in other non-human primate species, including macaques, and have exquisitely demonstrated alignment with human Tau [14], this has not yet been studied comprehensively in marmosets. Of the few studies in marmosets that have reported Tau pathology and phosphorylation sites [9–11], only one study investigated Tau isoform expression in a limited number of subjects [10]. Consequently, there remain several critical gaps concerning Tau expression in the marmoset and its relevance to AD. Specifically, age-dependent alterations of Tau isoform expression and phosphorylation residues related to pathological Tau aggregation and the subcellular distribution of Tau in the marmoset brain have yet to be comprehensively investigated.

Therefore, the present study aimed to investigate physiological 3R and 4R Tau expression comprehensively in brain tissues of unrelated marmosets across different age groups, subcellular distribution of Tau, as well as the aggregation-related phosphorylation residues and it is properties, which are critical foundational knowledge for understanding the relevance of marmoset model for the study of AD[1].

## 2.0 METHODS

### 2.1 Subjects

All experimental procedures involving animals were performed in accordance with state and federal laws, locally approved by The University of Pittsburgh Institutional Animal Care and Use Committee (IACUC), and were in line with and strictly adhered to the Guide for the Care and Use of Laboratory Animals [38].

#### 2.1.1 Marmosets

Outbred male and female common marmosets (*Callihrix jacchus*) were housed in an AAALAC-accredited facility at the University of Pittsburgh. Subjects were housed in pairs or family groups and maintained at a temperature range of 76–78 °F and 30–70% humidity, with a 12 h:12 h light/dark cycle (lights on at 7 am). Subjects were fed a diet consisting of twice daily provisions of commercial chow, including a purified diet, and supplemented with fresh fruit and vegetables daily with drinking water provided ad libitum. Foraging materials and enrichment were also provided daily. Subject demographics are provided in **Supplement Table 1**.

#### 2.1.2. Mice

Breeding pairs of C57BL/6J mice (JAX# 000664) were obtained from the Jackson Laboratory (Bar Harbor, ME) at 8-12 weeks of age. Mice were group-housed (n=2-4 per cage) with ad libitum food and water in a dedicated mouse housing room with a 12:12 light: dark cycle (lights on at 7 am). N=2 male offspring, aged 3 weeks and 17 months, were used for these studies. All experiments were conducted during the light cycle.

### 2.2 Tissue preparation

Brain tissues from unrelated outbred male and female marmosets and inbred C57BL/6J mice were obtained following humane euthanasia in accordance with the AVMA Guidelines for the euthanasia of animals (https://www.avma.org/resources-tools/avma-policies/avma-guidelines-euthanasia-animals). Briefly, mice (n=2) were anesthetized by isoflurane inhalant anesthesia (3-5% in O2) to the surgical plane of anesthesia, and the brain was dissected and flash-frozen following decapitation. Marmosets were sedated and anesthetized by intramuscular injection of ketamine (20–40 mg/kg) and intravenous injection of sodium pentobarbital (10–30 mg/ kg), respectively. Brain hemispheres were extracted following transcardial perfusion with ice-cold PBS for immunofluorescence staining, then fixed in 10% Neutral Buffered Formalin (NBF; Sigma Aldrich) until analysis. For biochemical analysis, brain hemispheres were rapidly extracted after perfusion and snap-frozen by isopentane with dry ice and stored at −80 °C until analysis. Tissue was collected within 12 hours of parturition from n=1 neonate found dead and immediately frozen and stored at -80 °C until analysis. Frozen inferior temporal cortex tissue of the brain from a de-identified Alzheimer’s Disease patient donor with confirmed Tau pathology was obtained from the Department of Pathology at the University of Pittsburgh, following CORID (Committee for Oversight of Research and Clinical Training Involving Decedents) approval and used as control tissue for western blot analysis. Postmortem frozen human brain of Alzheimer’s Disease (n=4) and non-demented control sections (n=4) of the frontal cortex were obtained from the Emory Alzheimer’s Disease Research Center brain bank (pathological traits described in **Supplemental Table 2)** and used for Mass spectrometry.

### 2.3 RNA isolation and reverse transcription cDNA

Prefrontal cortex tissue from n=3 marmosets (50-70mg), including a neonate, an adolescent (13 months), and an aged adult (9 years), were used for RNA extraction. Frozen brains were thawed on the wet ice, and total RNA was extracted by TRIzol Plus RNA purification kit (Invitrogen#12183555) according to the manufacturer’s protocol. On-column DNase treatment (PureLink™ DNase Set, ThermoFisher Scientific # 12185010) was performed to obtain DNA-free total RNA. The yields of the total RNA for each sample were determined using a Nanodrop one spectrophotometer (Thermo Fisher Scientific). 100 ng total RNA and oligo(dT)20 primer were used for reverse transcription by SuperScript™ III First-Strand Synthesis System (ThermoFisher Scientific #18080-051) according to the manufacturer-provided protocol to construct the cDNA library.

### 2.4 RT-PCR analysis and *MAPT* sequencing

A reverse transcribed cDNA library was used as a template to amplify *MAPT* isoforms (with and without exon10) using RT-PCR. The reaction was carried out by DreamTaq™ Hot Start Green DNA Polymerase (Thermo Fisher Scientific #MAN0015979). The PCR reaction was executed in ThermalCycler9 (Bio-Rad) under a touchdown PCR program (annealing Tm from 68°C to 63°C decrements of 0.5°C in every cycle for 10 cycles, followed by 63.5°C for 25cycles). The primer used in the reaction was designed based on marmoset 2N4R Tau mRNA sequence (NCBI database (MK630008): forward primer GTCAAGTCCAAGATCGGTTC; reverse primer TGGTCTGTCTTGGCTTTGGC. The PCR products were analyzed by electrophoresis on 4% (w/v) agarose gels. The amplified DNA products were labeled with GelGreen® Nucleic Acid Gel Stain dye (Biotium # 41004) and visualized by ChemiDoc Imaging Systems (Bio-rad, California, USA). Following visualization, each *MAPT* DNA band was excised from agarose gel, and DNA was purified by gel extraction kit (Qiagen# 2874 Maryland, USA) according to the manufacturer’s instructions. The purified DNA fragments were sequenced by Genewhiz (AZENTA life sciences, MA, USA) from both 5’ and 3’ terminals. The *MAPT* isoforms were confirmed using the Nucleotide BLAST program (NIH, National Library of Medicine).

### 2.5 Extraction of Sarkosyl soluble and insoluble Tau

The Sarkosyl soluble and insoluble fractions were extracted from the hippocampus, entorhinal cortex, and prefrontal cortex of marmoset frozen brain tissues and the frozen human AD brain cortex, similar to methods previously described [39]. Briefly, the brain tissue was homogenized using Potter-Elvehjem tissue homogenizer in nine volumes (wt/vol) ) of tris homogenize buffer:10mM Tris-HCl, pH7.4, 0.8M NaCl, 10% sucrose, 1mM EGTA, 2mM DTT(dithiothreitol), EDTA-free Pierce™ Protease Inhibitor Mini Tablet (Thermo fisher scientific# A32955), 1x phosphatase inhibitor cocktail I (Abcam, Waltham, MA #ab201112) with 0.1% Sarkosyl added, and centrifuged at 10,000 × g for 10 min at 4°C. Pellets were re-extracted using half the volume of the homogenization buffer, and resulting supernatants were pooled (S1). Additional Sarkosyl was added to the supernatant (S1) to reach a final concentration of 1% and rotated for an additional 1 hour at 4°C, followed by 60-min centrifugation at 300,000 × g at 4°C. The resulting pellets were resuspended in 100µL PBS as Sarkosyl-insoluble fraction (P2). The supernatants are Sarkosyl-soluble fraction (S2). Each fraction (S2, P2) was analyzed by western blotting.

### 2.6 Crude synaptosome fractionation

Crude synaptosome was isolated by sub-cellular fractionation as described[40]. Briefly, frozen marmoset prefrontal cortex and mouse brain tissues were homogenized using Potter-Elvehjem tissue homogenizer in nine volumes (w/v) of HEPES homogenization lysis buffer: 4mM HEPES, pH 7.4, 2mM EGTA, 0.32M sucrose, 2mM DTT with 1x Halt protease inhibitor cocktail (Thermo Fisher Scientific#78430), and 1x phosphatase inhibitor cocktail (Abcam, Waltham, MA #ab201112). The lysates were centrifuged at 1000 × g for 5 min at 4°C to remove nuclear material and cell debris. The supernatant (S1) was centrifuged at 12,000 × g for 15 min at 4°C, yielding supernatant (S2) and Pellets (P2). S2 was further centrifuged at 50,000 × g for 30 min at 4°C, yielding supernatant (S3), a cytosolic fraction. The P2, which is the crude synaptosome fraction, was resuspended in the HEPES lysis buffer with the addition of 0.1% TritonX100, and rotated for 1hour at 4°C followed by 60min centrifugation at 16,000 × g at 4°C which yielded supernatant (S4) as the extra-synaptic fraction and pellets (P4) as the postsynaptic density fraction (PSD). P4 was resuspended in HEPES lysis buffer with an additional 0.3% TritonX100. The fractions (S3, S4, P4) were analyzed by Western blotting.

### 2.7 Western Blot (WB)

For WB, the protein concentration of various fractions from different subjects was determined using a bicinchoninic acid (BCA) assay (ThermoFisher Scientific). Proteins were denatured by heating at 95°C for 5min, adding 1X Laemmli sample buffer with 2.5% of 2-Mercaptoethanol. Equivalent amounts of protein were loaded on 4-15% Mini-PROTEAN precast TGX gel (Bio-Rad). After electrophoresis, proteins were transferred to the nitrocellulose membrane using a turbo transfer system (Bio-Rad). The membranes were further blocked with EveryBlot blocking buffer (Bio-Rad #12010020) for 30min at room temperature and incubated with primary antibodies overnight at 4 °C. After washes (3X for 10min), the membranes were incubated for 1 hour in fluorescent-dye conjugated or HRP (horseradish peroxidase) conjugated anti-rabbit or anti-mouse secondary antibodies. The HRP-labeled membrane was further incubated with Pierce ECL substrate (Thermo Fisher Scientific# 32106). The target protein expression was visualized by ChemiDoc MP Imaging System (Bio-Rad, CA, USA). GAPDH was used as a loading control. For Sarkosyl insoluble fraction analysis, after visualization of AT8 targeted Tau protein, the membrane was stripped with 1Il Nitrocellulose stripping buffer (Thermo Fisher #J62541.AP) following the manufacturer’s instructions, then re-probed with RD3 and 4RTau antibodies. Primary antibodies used for western blotting in this study were: mouse anti-4R Tau monoclonal antibody-RD4 (1:800; Millipore sigma# 05-804); mouse anti-3R Tau monoclonal antibody-RD3 (1:800; Millipore sigma# 05-803); Rabbit anti-4RTau monoclonal antibody (1:1000; Cell signaling#79327); mouse anti-Tau monoclonal antibody-Tau5 (1:500; Thermo Fisher #AHB0042); mouse anti-human Tau monoclonal antibody-HT7(1:2000; Thermo Fisher # MN1000); mouse anti-S396/S404 phospho-Tau monoclonal antibody-PHF1 (1:500; gifted from the laboratory of Dr. Peter Davies, Department of Pathology, Albert Einstein College of Medicine, NY, USA); mouse anti-phospho-Tau(Ser202/Thr205) Monoclonal Antibody-AT8 (1:300; Thermo Fisher # MN1020); mouse anti-phospho-Tau (Thr231) monoclonal antibody-AT180(1:500; Thermo Fisher # MN1040); mouse anti-Tau oligomeric antibody-TOMA1(1:300; Thermo Fisher #MABN819); mouse anti-phospho-Tau (Thr181) monoclonal antibody-AT270 (1:1000; Thermo Fisher # MN1050); rabbit anti-phospho-Tau (Thr217) monoclonal antibody (1:1000; Cellsignaling # 51625); hFAB™ rhodamine conjugated anti-GAPDH antibody (1:2000; Bio-Rad #12004168); mouse anti-PSD95 monoclonal antibody(1:1000; abcam#ab2723); mouse anti-synaptophysin monoclonal antibody (1:1000; Thermo Fisher Scientific#MA1-213). Secondary antibodies used in this study were: HRP (horseradish peroxidase)-conjugated Goat-anti-mouse IgG secondary antibody (1:10000; Jackson ImmunoResearch # 115-035-003) and Alexa Fluor™ Plus 647 conjugated goat anti-rabbit IgG secondary antibody (1:3000; Thermo Fisher Scientific#A32733). For AD positive control, 0.3µL insoluble fraction was loaded, while for all marmoset samples, an equivalent volume (22.5µL) of insoluble fractions was loaded in each well.

### 2.7 Immunofluorescence (IF)

Perfused and fixed (10%NBF) marmoset brains were embedded in paraffin and sectioned into 4µm slices by microtome for fluorescent immunostaining. Briefly, the slices were de-paraffined and dehydrated, followed by citric-acid-based antigen retrieval (Vector Laboratories# H-3300) for 30 min in a steam cooker. After washing, the slices were incubated in 0.3% H2O2 at room temperature for 15min to block endogenous peroxidase activity. Following TBS wash (3x), the slices were incubated with 0.1% TritonX100 for 10min at room temperature, then blocked with 5% goat serum and 5% donkey serum in TBST (TBS with 0.1% TritonX100) at room temperature for 1 hour. After blocking, the slices were incubated with primary antibodies RD3 (1:200) and anti-4RTau (1:300) in a blocking medium at 4 °C overnight, followed by a 10 min wash (3x). Subsequently, the slices were incubated with secondary antibodies: Alexa Fluor 488-conjugated goat anti-rabbit Ab (Abcam #ab150077) and Alexa Fluor 647-conjugated donkey anti-mouse Ab (Abcam #ab150107) for 60 min at room temperature and mounted with ProLong Diamond Antifade Mounting Medium with DAPI (ThermoFisher #P36961). Slices without the incubation of the primary antibodies were used as controls. The images were taken with a Leica SP8 confocal microscope with a 20X objective lens with 3 times magnification.

### 2.8 Mass Spectrometry (MS)

Marmosets (n=7), mouse (n=1), and human (n=8) brain sections were analyzed by MS and illustrated in **Fig2A.** The frontal cortex of frozen brain sections from Human Alzheimer’s Disease patients and non-demented controls were obtained from the Emory Alzheimer’s Disease Research Center brain bank (pathological traits described in **Supplemental Table 2**). Marmoset, mouse, and human brain tissues were homogenized as described [41, 42]. Briefly, human brain, marmoset (prefrontal cortex), and mouse hemi brain were homogenized in Urea homogenization buffer (10 mM Tris, 100 mM NaH2PO4, 8M Urea, pH 8.5 with 1x Halt protease inhibitor (ThermoFisher) with ∼100 µL stainless-steel beads (0.9 to 2.0 mm, (NextAdvance) by a bullet blender at 4°C for 2 full 5-min intervals. Lysates were transferred to fresh tubes and sonicated three times at 30% amplitude on ice for 15s with 5s intervals. Lysates were centrifuged at 4°C for 5 min at 15,000 x g. Supernatants were collected and used for MS. The protein concentrations were determined through bicinchoninic acid assay (Pierce). The lysates were further reduced with 5mM DTT and alkylated with 10 mM iodoacetamide (IAA) at room temperature for 30min respectively, followed by diluting urea concentration to < 1M by adding 100 mM Tris-HCl, pH 8.0 buffer (v/v=9:1), and 1 mM CaCl2 buffer. To identify the Tau isoform-specific peptides, a parallelized dual digestion approach was executed with Trypsin and LysargiNase (LysArg); the latter cleaves at the N-terminus of K/R residues (**Fig 2B).** Briefly, 60µg of total protein per sample and 5µg recombinant Tau (rTau) (2N4RTau and 1N3R Tau) were digested with either Trypsin Protease (Pierce) or LysargiNase (EMD Millipore) at 1:50 w/w overnight at room temperature. The resulting peptides were acidified 1:9 (v/v) with acidification buffer (10% formic acid and 1% trifluoroacetic acid to quench enzyme activity and desalted by loading the peptides onto a 10 mg Oasis PRiME HLB 96-well plate (Waters), followed by washing twice with Buffer A (0.1% TFA) and then eluted with Buffer C (50% acetonitrile, ACN and 0.1% TFA). Desalted peptides were lyophilized with a CentriVap Centrifugal Vacuum Concentrator (Labconco) overnight. 3R-specific (KVQIVY) peptide generated by LysargiNase and 4R-specific (VQIINK) peptide generated by trypsin, which is shared in primary sequence across mice, marmosets, and humans were selected at the targeted Tau peptide sequences. A common Tau tryptic peptide shared across 3R and 4R Tau isoforms and species, SGYSSPGSPGTPGSR, to determine generalized equal loading across samples (**Fig 2B)**. Purified recombinant 3R and 4R Tau were used as positive controls to confirm peptide identification via unique MS/MS profiles. Recombinant 2N4R Tau was purchased from SignalChem, and it was maintained in the manufacturer storage buffer (50 mM Tris-HCl, pH 7.5, 150 mM NaCl, 0.25 mM DTT, 0.1 mM PMSF, 25% glycerol). 1N3R Tau was purchased from rPeptide and resuspended in 100 mM Tris-HCl pH 8.0 buffer (Invitrogen). 5 µg of total recombinant protein and 0.1 µg of enzyme were used per sample for digestion.

**Figure 1.**
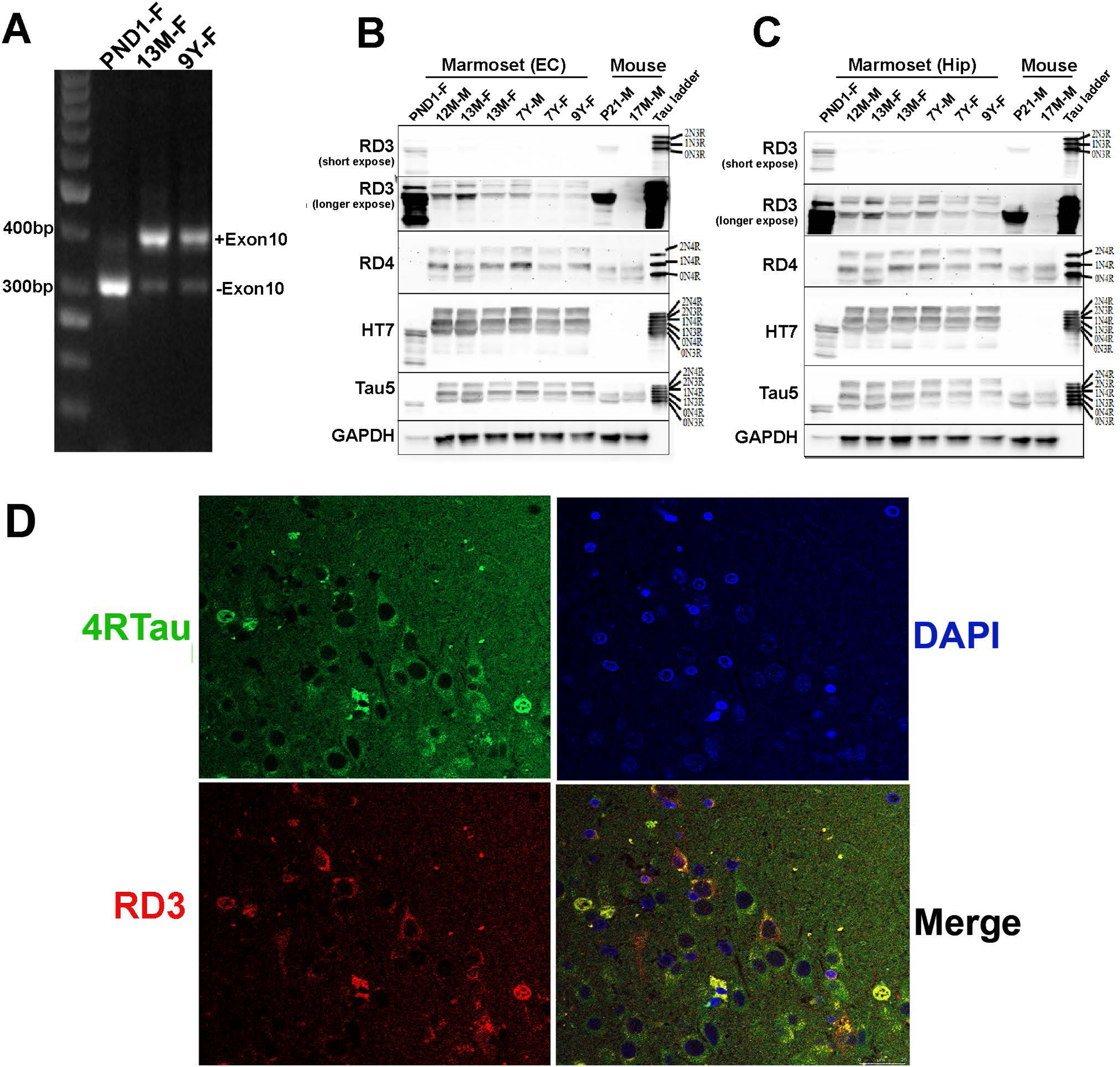
3R and 4R Tau expression in marmoset brain. **A**) RT-PCR products of marmoset frontal cortex. Subjects are three unrelated marmosets spanning an age range of postnatal day 1 through age 9 years. The PCR products without exon 10 (304bp) and with exon 10 MAPT (396bp) isoforms were amplified from the cDNA library of the prefrontal cortex. Lanes are identified as follows: Lane 1, DNA ladder: PND1 (postnatal day 1, female); 13M (13 months old, female); 9Y (9-year aged, female). Arrows indicating PCR product with or without exon10. **B)** 3R and 4R Tau isoform expression in marmoset Entorhinal cortex (EC), and **C)** Hippocampus (Hip) prepared as 1% Sarkosyl soluble lysate in individual outbred wild-type marmosets (postnatal day1 to 9 years old), and compared to mouse brain (PND 21 and 17 months old, respectively). Figure 1B and 1C: recombinant human Tau ladder showing 6 Tau isoforms. Row 1 is a short-time exposure to RD3 immunoblot, and row 2 is a longer exposure to RD3 immunoblot. 3R Tau was detected by anti-3R Tau specific antibody RD3, 4R Tau was detected by anti-4R Tau specific antibody RD4. Total Tau expression was determined by anti-human Tau specific antibody HT7 and anti-Tau antibody Tau5. GAPDH is a loading control. M: male; F: female; **D)** Immunofluorescence staining of 12-year aged male marmoset. Paraffin slides were stained with anti-3RTau (RD3) and anti-4RTau antibodies with Alex488 and Alex647 secondary antibodies, respectively. The red signals are 3RTau signals, and the green are 4RTau signals. Cell nuclei were stained with DAPI (blue). 3R/ 4R Tau co-localization in the merged image (orange). The scale bar is 25µm.

**Figure 2.**
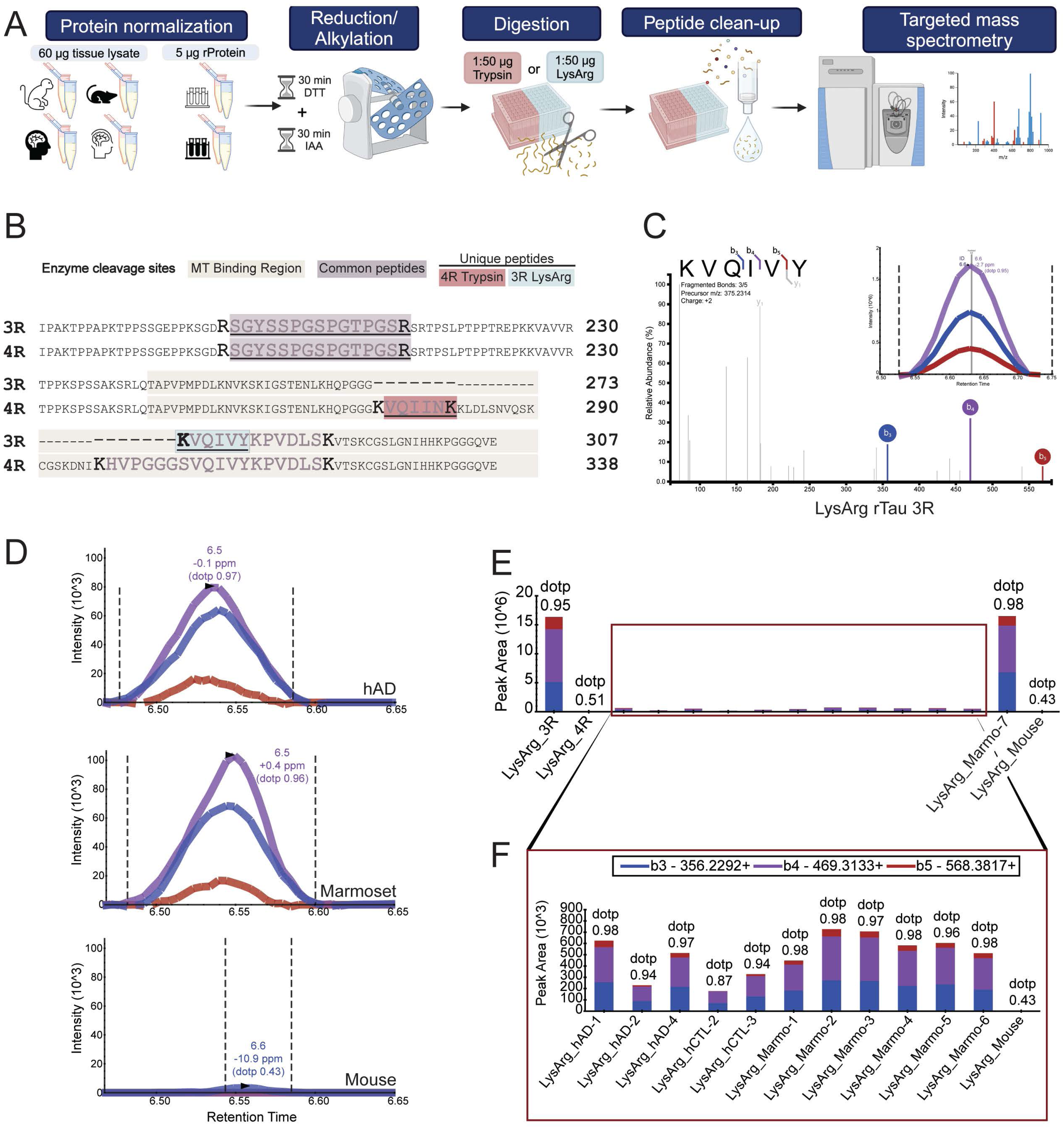
3R Tau expression in Marmosets brain by targeted mass spectrometry. **A)** Schematic mass spectrometry overview using a dual enzymatic digestion approach with Trypsin and LysargiNase (LysArg). Samples (n = 18 total) including 7 Marmosets, 4 human Alzheimer’s Disease (hAD) cases, 4 human non-demented control (hCTL), and 1 mouse were analyzed. **B)** Schematic diagram illustrating the primary sequence of human 3R and 4R Tau. Represented is the shared 3R and 4R peptide SGYSSPGSPGTPGSR (purple), unique 3R peptide, KVQIVY (blue), and unique 4R peptide, VQIINK (pink). **C)** MS/MS spectrum of the unique 3R Tau peptide, KVQIVY (m/z 375.2314, charge +2) generated by LysArg digestion of recombinant 3R Tau protein. The top three product ions (b3, b4, b5) are colored. The dot-product (dotp) value measures the similarity of the product ion pattern of the peptide to the recombinant 3R/4R Tau library, with a dotp value of 1 being a perfect match. The KVQIVY product ions for recombinant 3R Tau dotp value is 0.96. **D)** Peak area fragment ion intensities of the KVQIVY 3R specific peptide across species. The top panel displays human AD (dotp = 0.97), the middle panel presents marmoset (dotp = 0.96), and the bottom panel shows mouse (dotp = 0.43). The human and marmoset ion fragments and peak areas are comparable to those in the recombinant 3R Tau peptide. Additionally, retention time matching and overlapping fragment ions confirm the co-elution of the correct peptide in the targeted proteomics approach. **E)** Overview of the unique KVQIVY 3R peptide peak area product ion intensities across samples digested with LysArg. All samples underwent retention time window filtering to +/-15s of the predicted elution time determined by the SSRCalc 3.0 hydrophobicity retention time calculator. The best peak was selected for analysis, and the strongest intensities were displayed in rTau 3R and Marmo-7 (neonatal marmoset) samples, as expected. Recombinant 4R Tau (dotp = 0.51) and mouse (dotp = 0.43) samples do not display corresponding product ions as the 3R recombinant Tau. **F)** Inset of peak areas plot to allow visualization of biological samples on the same scale. All marmoset and human product ions collected (b3, b4, b5) are consistent with the recombinant 3R Tau (panel C).

Each sample was analyzed on a Q-Exactive HFX mass spectrometer (ThermoFisher Scientific) fitted with a Nanospray Flex ion source and coupled to an M-Class Acquity liquid chromatography system (Waters Corporation) essentially as described [43, 44]. The peptides were resuspended in 40 μL of loading buffer (0.1% TFA), and 1 µl was loaded onto a Waters CSH 1.7um C18 column (150um x 15 cm)). Elution was performed over a 10-min gradient at a nominal rate of 1500 nL/min with buffer B ranging from 1 to 20% (buffer A: 0.1% formic acid in water, buffer B: 0.1% formic acid in ACN) followed by a 5 min 99% wash.

The mass spectrometer was set to collect in PRM (parallel reaction monitoring) mode with an inclusion list consisting of each peptide (**Supplemental Table 3**). An additional full survey scan was collected to assess for possible interference. Full scans were collected at a resolution of 15,000 at 200 m/z with an automatic gain control (AGC) setting of 1× 105 ions and a max ion transfer (IT) time of 22ms. For PRM scans, the settings were: resolution of 30,000 at 200 m/z, AGC target of 1[×[106 ions, max injection time of 64ms, loop count of 4, MSX count of 1, isolation width of 1.6 m/z and isolation offset of 0.0 m/z. A pre-optimized normalized collision energy of 28% was used to obtain the maximal recovery of target product ions.

## 2.9 Data Analysis

### 2.9.1 RT-PCR and WB

The DNA bands were quantified using ImageJ (National Institute of Health). The relative ratio of MAPT with and without exon10 isoforms was derived from each band by dividing the optical density (OD) of interest from the same column. WB images were analyzed by ImageJ.

### 2.9.2 Mass spectrometry

#### 2.9.2.1 Spectral Library Generation

Data-dependent acquisition (DDA) LC-MS/MS for rTau samples were generated on an HFX Orbitrap essentially as described [44] and imported into Proteome Discoverer, PD (Thermo, version 2.5), using the basic consensus workflow and basic QE processing workflow with the addition of SequestHT and Percolator nodes. A background proteome database of 451 proteins and all Tau isoforms were incorporated for FDR correction. Only the input files and enzyme selection were adjusted between the Trypsin and LysargiNase library generations. Parameters were set to 2 maximum missed cleavages, 20 ppm precursor mass tolerance, and a 0.05 Da fragment mass tolerance. Variable modifications included methionine oxidation (+15.995 Da) and dynamic protein terminus modifications (N-term acetylation +42.011, met-loss –131.040, Met-loss+Acetyl –89.030). Carbamidomethyl +57.021 Da on cysteines was selected as a static modification. Percolator settings relied on concatenated validation based on q-value with a target false discovery rate (FDR) of 0.01 (strict) to 0.05 (relaxed).

#### 2.9.2.2 Skyline Product Ion and Peak Selection

The subsequent PD results files were imported into Skyline (MacCoss Lab, version 23.1.0.268) for downstream analysis. Skyline settings were as follows: either promiscuous LysN or trypsin enzyme, Human 2019 background proteome, and a minimum peptide length of 6. The peptide modifications were selected to match the PD parameters. The transition settings included the selection of the first to last product ions with precursor charges of 2 and 3; ion charges of 1 and 2; and ion types of y and b. The settings also included a 5 m/z precursor exclusion window and library match tolerance within 0.1 m/z.

The most intense ions from the library spectrum were selected. The SSRCalc 3.0 (300A) hydrophobicity retention time calculator was used for all peptides, and the peptide peak selection was limited to a ± 15-second time window of this predicted elution time (**Supplemental Table 4)** and within ±5 ppm of the library spectrum to ensure confidence in the resulting product ion peaks. Peaks were manually inspected for discordant matches, and if identified sample peaks did not fall into the predicted retention time range, they were excluded from the analysis. Human AD3, CTL1 (control 1), and CTL4 (control4) were excluded from the analysis as these did not meet the retention time filtering. A Ratio Dot Product (dotp) value was used to describe the similarity of the experimental spectra to the comparative reference recombinant protein spectra, with a dotp value of 1.0 denoting the highest similarity. All PRM raw data files, including library and Skyline analysis files, are available on Synapse (syn52356795 and syn52895027).

#### 2.9.2.3 Base Peak Filtering and Resulting Chromatograms

Recombinant rTau. RAW files were loaded into the XCalibur Qual Browser application (Thermo Xcalibur version 4.2.47, January 24, 2019). The precursor m/z from Skyline was selected for base peak filtering. The spectrum list with m/z and intensity values (**Supplemental Table 5**) was exported and uploaded into the interactive peptide spectral annotator [45] to generate chromatograms.

## 3.0 RESULTS

### 3.1 *MAPT* mRNA splicing in marmoset brain

Alternative splicing of exon 10 of *MAPT* results in 3R and 4R Tau isoforms in the human brain. The inclusion or exclusion of exon 10 gives rise to 4-repeat (4R) and 3-repeat (3R) tau, respectively [13]. To determine whether the 3R and 4R Tau isoform splicing occurs in marmosets, total mRNA from the prefrontal cortex of unrelated marmosets across different ages from neonate through 9 years was extracted and transcribed to the cDNA library. The exon 10 splicing was determined by RT-PCR using a cDNA library as a template from each sample. As illustrated in **Figure 1A**, *MAPT* without exon 10 mRNA was predominately detected in neonatal marmoset brains relative to *MAPT* with exon 10 (**Fig.1A**, lane 1). In contrast, *MAPT* mRNA with exon 10 (**Fig. 1A**, top band of lane 2, 3) was predominately detected in brains from the 13-month and 9-year-aged marmosets, relative to *MAPT* without exon 10 (**Fig. 1A**, bottom band of lane 2, 3). The sequence of each fragment was confirmed by Sanger DNA sequencing.

The 304bp fragment without exon 10 was 100% aligned with 0N3R *MAPT* mRNA of marmoset (MGenBank: MK630010.1), and the 397bp fragment with exon 10 was 100% matches with 0N4R *MAPT* mRNA of marmoset (MGenBank: MK630009.1). By qPCR, 3R Tau mRNA quantification in brain tissues of marmosets of different ages also demonstrated 3R Tau mRNA expression in the neonate with approximately 10-15-fold higher expression relative to 3R Tau mRNA expression in the adolescent and the adult marmoset brains (**Supplemental Figure S1**). These results confirm that exon 10 splicing results in 3R and 4R Tau isoforms in marmoset brain throughout the marmoset lifespan, with 3R Tau predominantly expressed in neonatal marmoset brain and 4R Tau predominantly expressed in adolescent/adult marmoset brain.

### 3.2 3R and 4R Tau protein expression in marmoset brain

To further verify 3R and 4R Tau protein expression, the Sarkosyl soluble fraction from the hippocampus (Hip) and entorhinal cortex (EC) were evaluated in n=7 marmosets of various ages and analyzed by WB. As presented in **Figure 1B-C**, in the neonatal marmoset brain, 3R Tau was predominantly expressed relative to 4R Tau in both the EC (**Fig. 1B**) and Hip (**Fig. 1C**), which was also observed as expected in postnatal mouse brain (PND 21), but not in adult mouse brain (17 months of age). In contrast, 4R Tau was predominantly expressed in adolescent marmoset brain regions (**Figure 1B-C**, 12-13 months) and adult marmoset brain regions (**Figure 1B-C**, 7-9 years) relative to 3R Tau expression, which was expressed as expected in both postnatal mouse brain and adult mouse brain regions. The total Tau expression was detected by HT7 antibody (human Tau-specific antibody) and Tau5 antibody. HT7 antibody only binds to marmoset Tau, whereas Tau5 binds to both marmoset and mouse Tau, demonstrating the differential isoform expression pattern of Tau in marmoset relative to the mouse. To further confirm the expression of Tau isoforms in marmoset, immunofluorescent staining (IF) and immunohistochemistry (IHC) were performed. As presented in **Figure 1D**, 4R Tau (green) and 3R Tau (red) signals were observed in the hippocampus region of a 12-year-old male marmoset. 4R Tau signal was strongest in the neuronal soma and proximal/distal segments of apical dendrites. 3R Tau shows puncta-like signals co-expressed with 4R Tau in neuronal soma (**Fig. 1D**, merge). No 3R and 4R tau positive signals were observed in slices incubated without the primary antibodies (**Supplemental Figure S3**). Immunohistochemistry (IHC) re-confirmed the observation of IF within the same subject (**Supplemental Figure S4**). Of note is that the 3R and 4R tau expression patterns were similar to those observed in IF. These data are consistent with RT-PCR data, confirming the presence of both 3R and 4R Tau isoforms in the marmoset brain across the lifespan.

### 3.3. Mass spectrometry of 3R and 4R Tau

For further validation of 3R and 4R Tau isoforms in the marmoset brain, a highly specific and sensitive targeted parallel reaction monitoring (PRM) mass spectrometry assay was implemented. Of note, the KVQIVY peptide was found in the recombinant 3R tau with a near-perfect match to the MS/MS spectrum library (dotp = 0.95) displaying b3, b4, and b5 product ions (**Fig. 2C**). As expected, the human ion fragments and peak areas were equivalent to those in the recombinant 3R tau library with identified peaks ≤0.7 ppm and an average retention time (RT) of 6.53 min (**Fig, 2D**, top graph). The pattern of the marmoset product ions mirrored the human results with identified peaks less than ≤0.6 ppm and an average RT of 6.53 (**Fig. 2D**, middle graph). In contrast, in the mouse brain lysate, we identified a peak at -10.9 ppm, which was outside of the mass accuracy threshold (±5 ppm) and did not correspond to the product ions of the 3R library or additional biological samples with a poor dotp of 0.43 (**Fig. 2D**, bottom panel). As expected, neither the 4R rTau isoform nor the mouse samples matched the 3R rTau library spectrum. Dotp values were 0.51 and 0.43, respectively (**Fig. 2E**, sample 2 and 15). The 3R rTau isoform (sample1) (dotp = 0.95, RT = 6.65) and neonatal marmoset (sample14) (dotp = 0.98, RT = 6.58) displayed KVQIVY peptide peak areas of 1x10^6^, which was a 10-20-fold higher over other marmoset and human samples, suggesting neonatal marmoset have highest 3R Tau isoform abundance (**Fig. 2E**). The human and marmoset samples highly matched the 3R rTau library MS/MS spectra. The average dotp values of human samples were 0.94, while marmoset samples were 0.98 (**Fig. 2F**). The shared 3R/4R Tau peptide, SGYSSPGSPGTPGSR, was included as a Tau control and displayed generally equivalent loading across biological samples (**Fig 3A**) with abundant y7, y10, and y11 product ions (**Fig. 3B**). The 4R-specific peptide, VQIINK, was also included to assay the 4R Tau isoform. All biological samples, mouse (dotp = 0.99), human (avg dotp = 0.98), and marmoset (avg dotp = 0.98), highly matched the 4R rTau library spectra (**Fig. 3C**). Abundant y3, y4, and y5 product ions were represented in the recombinant MS/MS spectrum of tryptic 4R rTau (**Fig. 3D**). Taken together, these data show that humans and marmosets express both 3R and 4R Tau isoforms into adulthood, and neonatal marmoset predominantly express 3R Tau with trace levels of 4R Tau.

**Figure 3.**
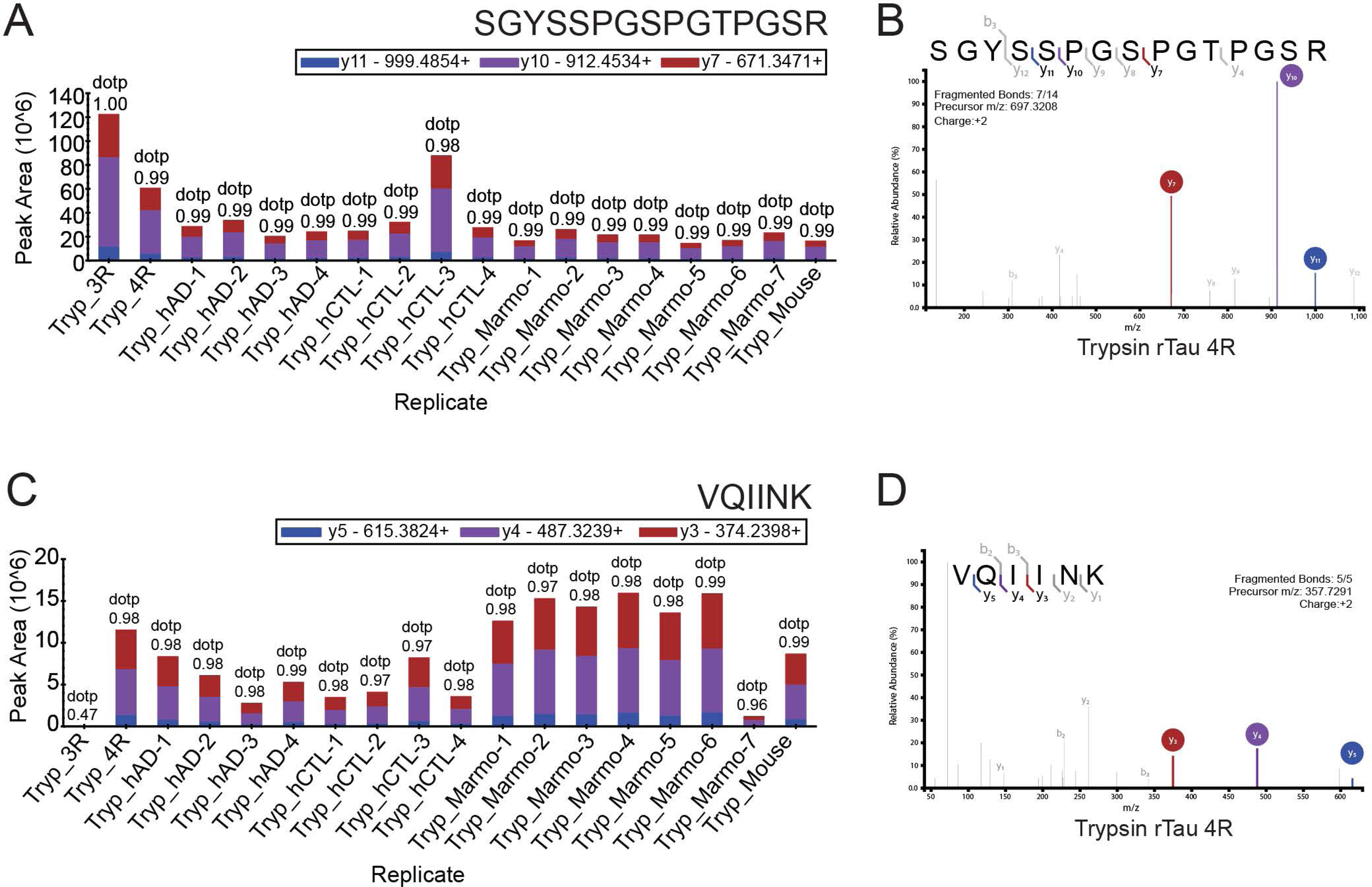
Total tau and 4R Tau expression in Marmosets by targeted mass spectrometry. **A)** Overview of the peak area product ion intensities of the tryptic 3R and 4R shared Tau peptide, SGYSSPGSPGTPGSR, providing evidence of Tau protein abundance across all samples. **B)** MS/MS spectrum of the 3R and 4R Tau shared peptide, SGYSSPGSPGTPGSR (m/z 697.3208, charge +2) generated by tryptic digestion of 4R recombinant Tau. The top three product ions (y7(red), y10 (purple), y11(blue)) are colored. **C)** Overview of the peak area product ions of the 4R unique tryptic peptide, VQIINK. The fragment ions were consistent with the 4R Tau library across all samples except for the recombinant 3R Tau (dotp = 0.47). **D)** Representative MS/MS spectrum of Tau 4R specific, VQIINK (357.7291 m/z, charge +2) peptide generated by tryptic digestion of recombinant 4R Tau. The top three ion products (y3(red), y4(purple), y5(blue)) are colored. Samples (n = 18 total) including 7 Marmosets, 4 human Alzheimer’s Disease (hAD) cases, 4 human non-demented control (hCTL), and 1 mouse were analyzed. Tryp-3R was recombinant human 3R Tau, and Tryp-4R was recombinant human 4RTau.

### 3.4 Tau isoform expression in the synaptic region

Increasing evidence indicates that Tau localizes in synaptic subcellular regions and plays essential roles in synaptic function [24–26]. To identify whether 3R Tau and 4R Tau were localized in synaptic subregions of adult marmoset neurons, a crude synaptosome was extracted from the prefrontal cortex, then further fractionated to extra-synaptic and postsynaptic density fractions, and analyzed by WB. As illustrated in **Figure 4**, 3R and 4R Tau were observed in the synaptic region (fraction S4, P4) along with predominant 4R Tau from three unrelated adult marmosets (ages 7 to 9 years). 3R Tau was primarily expressed in the cytosolic fraction, and the synaptic Tau was mainly distributed in the non-PSD synaptic region, albeit with individual variation. As expected, no 3R Tau isoform was detected in mouse brain extracts (**Fig. 4**). These data confirm the expression of both 3R and 4R Tau isoforms in synaptic regions of the marmoset brain.

**Figure 4.**
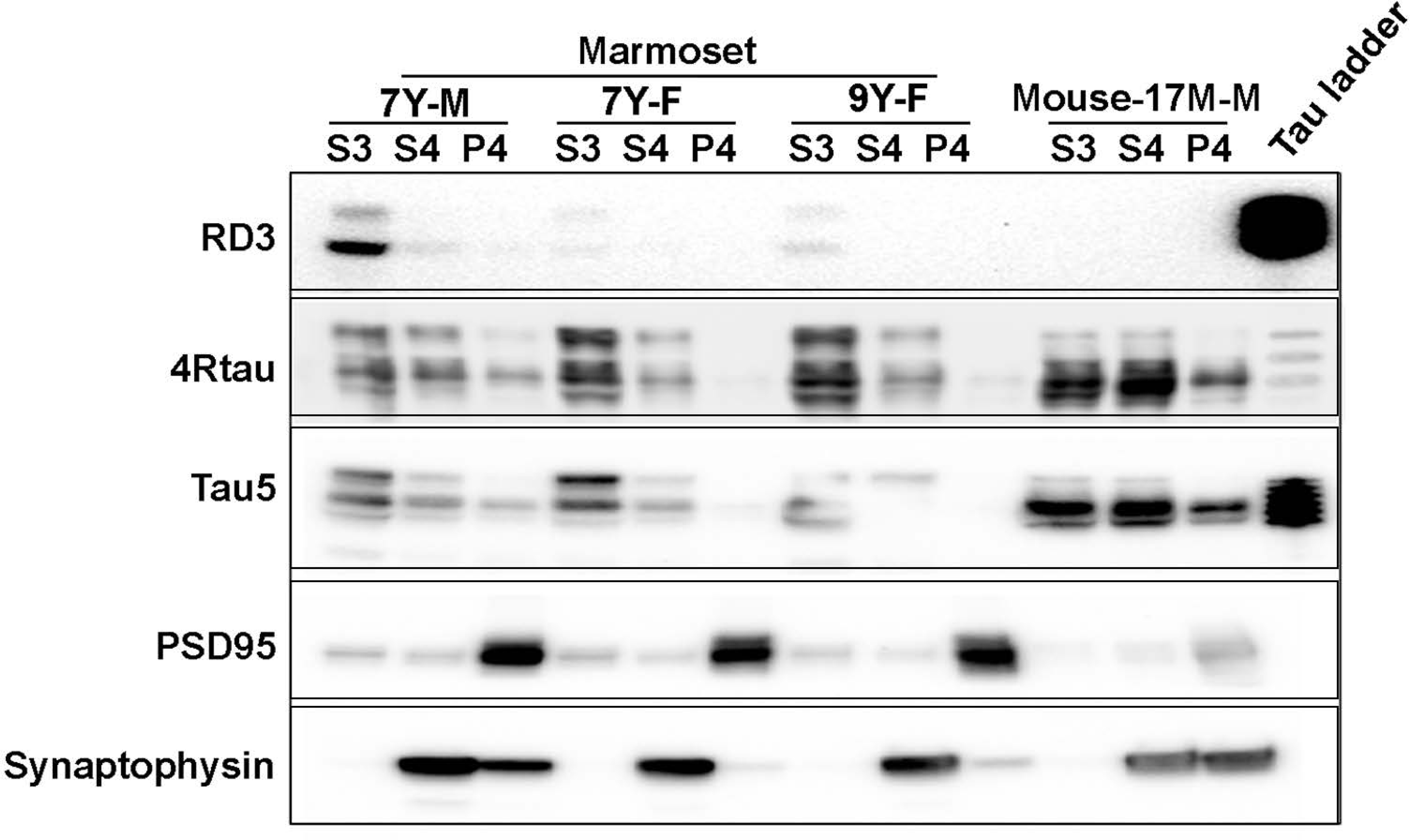
4R Tau and 3R Tau were expressed in the synaptic region of the marmoset prefrontal cortex. The prefrontal cortex of n=3 unrelated adult marmosets were fractionated, and 3R Tau and 4R Tau expression were determined by RD3 and 4R Tau antibodies. Adult C57BL/6J mouse brain was used as negative control for 3R Tau. The right lane is the recombinant human Tau ladder. S3: cytosolic fraction; S4: Extra-synaptic fraction, determined by an anti-synaptophysin antibody; P4: postsynaptic density fraction, determined by the anti-PSD95 antibody. Total Tau was determined by the anti-Tau5 antibody. M: male; F: female. 7 years (7Y); 9 years (9Y).

### 3.4 Tau phosphorylation, oligomerization in normal marmoset brain

Tau phosphorylation has been associated with Tau misfolding, accumulation, and formation of AD-like pathology [46–48]. To evaluate whether similar Tau phosphorylation sites are present in the marmoset brain, the Sarkosyl soluble fractions were analyzed by WB. As presented in **Figure 5**, Tau phosphorylation sites (T181, T231, T217) were phosphorylated in the soluble fraction of adolescent and adult marmoset brain (**Fig. 5A**) and present with oligomer-like properties, as detected by the Tau oligomer-specific monoclonal antibody TOMA1. Similar properties were observed in AD brain extract (**Fig. 5A**). To identify the possibility of the formation of high molecular weight Tau aggregates, the Sarkosyl insoluble fractions were analyzed by WB, with PHF1 and AT8 antibodies (**Fig. 5B**). The 1% Sarkosyl insoluble Tau in hippocampal extracts from n=3 adult marmosets were phosphorylated at S396/S404, in the absence of high molecular weight aggregates but not in neonatal and adolescent marmoset brains (**Fig. 5B**, top panel). A faint 200kD high molecular weight AT8 positive Tau was detected in adult hippocampus extraction but not in neonatal and adolescent marmoset brains (**Fig. 5B**). To confirm the isoform expression in AT8 positive aggregates, the membrane probed with AT8 was stripped and re-probed with anti-3R Tau and anti-4R Tau antibodies. Smeared 4R Tau bands (red) were observed in all samples, including human AD brain extracts, except for the neonatal sample, and two intense bands at 55kd and 60kd were observed in the brains from the adolescent (12 months) and the adult (7 years) marmosets. RD3 (green) was faintly stained with a smeared band in all samples, with the 42kd 3RTau band distinctly observed in the neonatal sample (**Fig. 5B**). These data indicated that T181, T231, T217, S202/T205, and S396/S404 phosphorylation sites were detected in the marmoset brain, with evidence of forming high molecular weight aggregates. Additional studies are required to further confirm the phosphorylation and aggregation properties of marmoset Tau.

**Figure 5.**
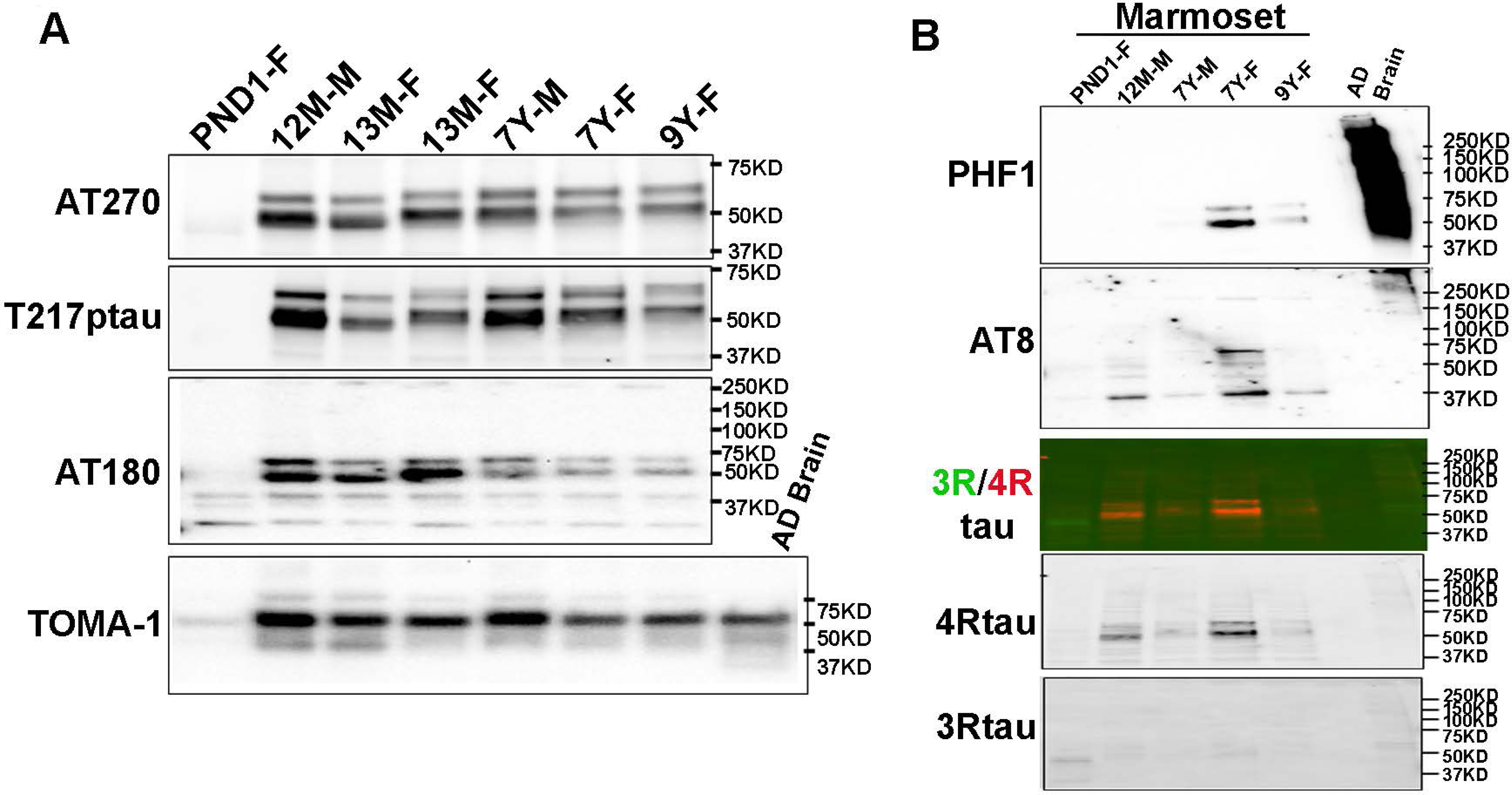
Phosphorylated Tau in Sarkosyl soluble and insoluble fractions from marmoset brain. Adult marmoset Hippocampus (Hip) was extracted as 1% Sarkosyl soluble and insoluble fractions in individual outbred unrelated marmosets and analyzed via western blots. **A)** The soluble extract of n=7 marmosets. The phosphorylation sites were determined by AT270, an anti-T181phospho-Tau monoclonal antibody; T217pTau, an anti-T217 phospho-Tau antibody; and AT180: anti-T231 phospho-Tau antibody. The Tau oligomer in the soluble fraction was confirmed by antibody TOMA1, an anti-human oligomeric Tau monoclonal antibody. **B)** Hyperphosphorylated high molecular weight Tau aggregates were detected in Sarkosyl insoluble fraction of adult marmoset hippocampus by AD Tau specific antibody PHF1 and AT8. RD3 and anti-4RTau antibodies confirmed the presence of 3R and 4R Tau in the Tau aggregates. Lanes 1 to 5 are marmoset samples from different ages and sexes. Lane 6 (blank); Lane 7, AD patient inferior temporal cortex extraction. Postnatal day 1 (PND1); 12 months (12M); 7 years (7Y); 9 years (9Y); male (M); female (F).

## 4.0 DISCUSSION

The present study is the first to confirm the expression of both 3R and 4R Tau isoforms in the marmoset brain throughout the lifespan. These results emphasize the relevance of the marmoset as a model system for the study of AD and related dementias[1].

Few studies have evaluated Tau expression and localization in the marmoset brain, and only one study focused on Tau isoform expression [9–12]. In contrast to that single report that marmosets do not express the 3R Tau isoform as adults, which was limited to only analysis of two marmosets [10] and may also be related to lower limits of detection of the reagents used, our comprehensive evaluation of Tau isoform expression in eight unrelated individual subjects across age ranges using multiple methodologies inclusive of RT-PCR, DNA fragments sequencing, western blot, immunohistochemistry, mass spectrometry, synaptosome fractions, all consistently verify the presence of both 3R and 4R Tau isoforms in marmoset brain.

The main difference between the previous report and our present results may be related to the detection methods and reagents used. Specifically, in the present study, we not only observed the DNA bands from RT-PCR products corresponding to mRNA of marmoset 3R Tau and 4R Tau but also amplified and sequenced their DNA fragments, which were matched with published marmoset *MAPT* mRNA isoforms from the MGenbank database (MK630010.1 and MK630009.1)[10]. Furthermore, we examined Tau isoforms in the Sarkosyl soluble and insoluble fractions, which is a standard protocol used to extract pathological Tau from human AD brain tissue [39], and immunofluorescent staining (IF) and IHC to visualize the 3R and 4R Tau in brain tissue (**Fig. 1D**). In addition, we performed a precise targeted mass spectrometry analysis via parallel reaction monitoring to confirm and extend these observations, including identifying 3R and 4R peptide sequences that correspond with those peptides unique to human. It is important to note that even with these robust methods, the presence of pathological Tau aggregates in these marmoset brain samples was scarce[4], especially relative to what is typically observed in the brains of AD patients[49–51]. This discrepancy may very well be related to the marmoset age at the time of death in the present set of experiments, as well as their limited longevity in captivity, as other laboratories have also reported only mild presentations of pathological Tau in the marmoset brain along with significant variability across individuals [9, 11]. In addition to IF results, we also observed AT8 and PHF1 positive aggregates in the Sarkosyl insoluble fraction along with 3R and 4RTau isoform expression **(Fig. 5)**. These Tau aggregates were observed only in older adult marmoset samples and absent in the adolescent samples. These findings may implicate an early stage of Tau aggregation. Relatedly, the present observation of 3R and 4R Tau expression in synaptosomes provides insight into potential trans-synaptic propagation, which is widely recognized as a consequence of Tau pathology trans-synaptic propagation and subsequent synaptic dysfunction in AD [31, 52]. Ongoing studies in our laboratory using both genetic and Tau seeding approaches will ultimately help to understand whether the marmoset is susceptible to Tau aggregation, propagation, and significant spreading, which may otherwise be attenuated due to their short lifespan in captivity relative to other non-human primate species which have more extensive NFT presentation [11, 53–56].

Extending our observation of the presence of both 3R and 4R Tau isoforms in marmoset brain, we sought to understand if marmosets recapitulate similar 4R/3R Tau ratios as described in human brain that vary across healthy and pathological conditions. Alternative *MAPT* exon 10 splicing is a complex process regulated by short cis-elements present both in exon 10 and in introns 9 and 10. In humans, 3R Tau is predominantly expressed in fetal and neonatal brains, while 3R and 4R Tau are expressed in the adult brain in roughly equal proportions[13, 14]. While the *MAPT* sequences of exon 10 and the stem-loop of intron10 are conserved between humans and marmosets[10, 20], and the predominant expression of 3R relative to 4R in the neonatal marmoset brain is conserved with the human neonatal 3R/4R ratio[16], the 3R/4R ratio in adolescent and adult marmosets was different, with 4R predominantly expressed with roughly a 5:1 ratio relative to 3R of the samples analyzed from mRNA in the present study (**Fig. 1A** and **Supplemental Figure 2**). This may indeed be the normative physiological expression pattern of 3R/4R Tau in healthy aging marmosets, though further detailed quantification will be required to confirm the translation and transcription levels of 3R and 4R Tau isoforms across colonies, as well as to understand if specific pathological conditions may result in a divergent ratio. Relatedly, this may also provide an additional explanation for the lack of observation of the 3R Tau isoform in adult marmosets, as previously reported [10]. Marmoset populations are genetically diverse and typically maintained in the laboratory as outbred colonies. Given that genetics may play a role in Tau isoform expression, at least in humans, it is possible that whole genome sequencing (WGS) data can also reveal insights into the differences across studies. However, WGS data have not yet been reported.

Tau hyperphosphorylation is associated with functional changes related to pathological conditions[25, 57, 58]. Phosphorylation in several residues contributes to the formation of pathological Tau aggregates in the human AD brain, notably at T217, T181, and T231, which have also been demonstrated as biomarkers of AD in tissues and fluids [35, 59]. In the present study, we demonstrate that these phosphorylation sites are naturally phosphorylated in the hippocampus of adolescent and adult marmosets. These results confirm and extend previous immunohistochemistry reports of hyperphosphorylated T231 Tau in adolescent and aged marmoset brains[9–11]. Furthermore, we also confirmed with AT8 and PHF1 epitopes the phosphorylation in Sarkosyl-insoluble fractions in adult marmoset hippocampus, which reproduces previous findings of AT8 expression in aged marmoset brain by immunohistochemistry[11].

Taken together, the present results confirm the expression of Tau isoforms in the marmoset brain and provide important and novel findings on tau homeostasis under physiological conditions. While further research is necessary to thoroughly investigate the role of pathological Tau formation in the marmoset and the trajectory of tau neurotoxicity that leads to neurodegenerative diseases, these findings highlight the importance of the marmoset as a model system for studying primate-specific mechanisms of AD, including its utility for evaluating interventions aimed at stopping or preventing AD.

## ACKNOWLEDGEMENTS

The authors are grateful to our dedicated marmoset veterinary and husbandry colleagues who provide exceptional care of the marmosets and assist with these studies. Mouse anti-S396/S404 phospho-Tau monoclonal antibody-PHF1 was gifted from the laboratory of Dr. Peter Davies, Department of Pathology, Albert Einstein College of Medicine, NY, USA and provided by Drs. Philippe Marambaud and Jeremy Koppel through the Feinstein Institutes for Medical Research. As part of the NIA funded Open Science Initiative, all data and protocols are made available through the AD Knowledge Portal (https://adknowledgeportal.synapse.org/Explore/Programs/DetailsPage?Program=MARMO-AD).

## Human subject consent

De-identified tissue was provided through approved resources at the University of Pittsburgh and Emory University that are exempt.

## REFERENCES

[1] Philippens I, Langermans JAM. Preclinical Marmoset Model for Targeting Chronic Inflammation as a Strategy to Prevent Alzheimer’s Disease. Vaccines (Basel). 2021;9.

[2] Rothwell ES, Freire-Cobo C, Varghese M, Edwards M, Janssen WGM, Hof PR, et al. The marmoset as an important primate model for longitudinal studies of neurocognitive aging. Am J Primatol. 2021;83:e23271.

[3] Sasaguri H, Hashimoto S, Watamura N, Sato K, Takamura R, Nagata K, et al. Recent Advances in the Modeling of Alzheimer’s Disease. Front Neurosci. 2022;16:807473.

[4] Perez-Cruz C, Rodriguez-Callejas JD. The common marmoset as a model of neurodegeneration. Trends Neurosci. 2023;46:394–409.

[5] Sukoff Rizzo SJ, Homanics G, Schaeffer DJ, Schaeffer L, Park JE, Oluoch J, et al. Bridging the rodent to human translational gap: Marmosets as model systems for the study of Alzheimer’s disease. Alzheimers Dement (N Y). 2023;9:e12417.

[6] Hayashi T, Hou Y, Glasser MF, Autio JA, Knoblauch K, Inoue-Murayama M, et al. The nonhuman primate neuroimaging and neuroanatomy project. Neuroimage. 2021;229:117726.

[7] Fukushima M, Ichinohe N, Okano H. Neuroanatomy of the Marmoset. In: Marini R, Wachtman L, Tardif S, Mansfield K, Fox J, editors. The Common Marmoset in Captivity and Biomedical Research: Academic Press; 2019. p. 43-62.

[8] Geula C, Nagykery N, Wu CK. Amyloid-beta deposits in the cerebral cortex of the aged common marmoset (Callithrix jacchus): incidence and chemical composition. Acta Neuropathol. 2002;103:48–58.

[9] Rodriguez-Callejas JD, Fuchs E, Perez-Cruz C. Evidence of Tau Hyperphosphorylation and Dystrophic Microglia in the Common Marmoset. Front Aging Neurosci. 2016;8:315.

[10] Sharma G, Huo A, Kimura T, Shiozawa S, Kobayashi R, Sahara N, et al. Tau isoform expression and phosphorylation in marmoset brains. J Biol Chem. 2019;294:11433–44.

[11] Arnsten AFT, Datta D, Preuss TM. Studies of aging nonhuman primates illuminate the etiology of early-stage Alzheimer’s-like neuropathology: An evolutionary perspective. Am J Primatol. 2021;83:e23254.

[12] Rodriguez-Callejas JD, Fuchs E, Perez-Cruz C. Atrophic astrocytes in aged marmosets present tau hyperphosphorylation, RNA oxidation, and DNA fragmentation. Neurobiol Aging. 2023;129:121–36.

[13] Goedert M, Spillantini MG, Potier MC, Ulrich J, Crowther RA. Cloning and sequencing of the cDNA encoding an isoform of microtubule-associated protein tau containing four tandem repeats: differential expression of tau protein mRNAs in human brain. EMBO J. 1989;8:393–9.

[14] Goedert M, Jakes R. Expression of separate isoforms of human tau protein: correlation with the tau pattern in brain and effects on tubulin polymerization. EMBO J. 1990;9:4225–30.

[15] Hefti MM, Farrell K, Kim S, Bowles KR, Fowkes ME, Raj T, et al. High-resolution temporal and regional mapping of MAPT expression and splicing in human brain development. PLoS One. 2018;13:e0195771.

[16] Kosik KS, Orecchio LD, Bakalis S, Neve RL. Developmentally regulated expression of specific tau sequences. Neuron. 1989;2:1389–97.

[17] Niblock M, Gallo JM. Tau alternative splicing in familial and sporadic tauopathies. Biochem Soc Trans. 2012;40:677–80.

[18] Glatz DC, Rujescu D, Tang Y, Berendt FJ, Hartmann AM, Faltraco F, et al. The alternative splicing of tau exon 10 and its regulatory proteins CLK2 and TRA2-BETA1 changes in sporadic Alzheimer’s disease. J Neurochem. 2006;96:635–44.

[19] Park SA, Ahn SI, Gallo JM. Tau mis-splicing in the pathogenesis of neurodegenerative disorders. BMB Rep. 2016;49:405–13.

[20] Liu F, Gong CX. Tau exon 10 alternative splicing and tauopathies. Mol Neurodegener. 2008;3:8.

[21] Tai HC, Serrano-Pozo A, Hashimoto T, Frosch MP, Spires-Jones TL, Hyman BT. The synaptic accumulation of hyperphosphorylated tau oligomers in Alzheimer disease is associated with dysfunction of the ubiquitin-proteasome system. Am J Pathol. 2012;181:1426–35.

[22] Chen Q, Zhou Z, Zhang L, Wang Y, Zhang YW, Zhong M, et al. Tau protein is involved in morphological plasticity in hippocampal neurons in response to BDNF. Neurochem Int. 2012;60:233–42.

[23] Wang Y, Mandelkow E. Tau in physiology and pathology. Nat Rev Neurosci. 2016;17:5–21.

[24] Hanger DP, Goniotaki D, Noble W. Synaptic Localisation of Tau. Adv Exp Med Biol. 2019;1184:105–12.

[25] Wang Y, Mandelkow E. Tau in physiology and pathology. Nature Reviews Neuroscience. 2016;17:22–35.

[26] Kimura T, Whitcomb DJ, Jo J, Regan P, Piers T, Heo S, et al. Microtubule-associated protein tau is essential for long-term depression in the hippocampus. Philos Trans R Soc Lond B Biol Sci. 2014;369:20130144.

[27] Mills F, Bartlett TE, Dissing-Olesen L, Wisniewska MB, Kuznicki J, Macvicar BA, et al. Cognitive flexibility and long-term depression (LTD) are impaired following beta-catenin stabilization in vivo. Proc Natl Acad Sci U S A. 2014;111:8631–6.

[28] Fa M, Puzzo D, Piacentini R, Staniszewski A, Zhang H, Baltrons MA, et al. Extracellular Tau Oligomers Produce An Immediate Impairment of LTP and Memory. Sci Rep. 2016;6:19393.

[29] Maeda S, Sahara N, Saito Y, Murayama S, Ikai A, Takashima A. Increased levels of granular tau oligomers: an early sign of brain aging and Alzheimer’s disease. Neurosci Res. 2006;54:197–201.

[30] Polydoro M, Dzhala VI, Pooler AM, Nicholls SB, McKinney AP, Sanchez L, et al. Soluble pathological tau in the entorhinal cortex leads to presynaptic deficits in an early Alzheimer’s disease model. Acta Neuropathol. 2014;127:257–70.

[31] Frost B, Jacks RL, Diamond MI. Propagation of tau misfolding from the outside to the inside of a cell. J Biol Chem. 2009;284:12845–52.

[32] Kaniyappan S, Chandupatla RR, Mandelkow EM, Mandelkow E. Extracellular low-n oligomers of tau cause selective synaptotoxicity without affecting cell viability. Alzheimers Dement. 2017;13:1270–91.

[33] Wesseling H, Mair W, Kumar M, Schlaffner CN, Tang S, Beerepoot P, et al. Tau PTM Profiles Identify Patient Heterogeneity and Stages of Alzheimer’s Disease. Cell. 2020;183:1699–713 e13.

[34] Tepper K, Biernat J, Kumar S, Wegmann S, Timm T, Hubschmann S, et al. Oligomer formation of tau protein hyperphosphorylated in cells. J Biol Chem. 2014;289:34389–407.

[35] Suarez-Calvet M, Karikari TK, Ashton NJ, Lantero Rodriguez J, Mila-Aloma M, Gispert JD, et al. Novel tau biomarkers phosphorylated at T181, T217 or T231 rise in the initial stages of the preclinical Alzheimer’s continuum when only subtle changes in Abeta pathology are detected. EMBO Mol Med. 2020;12:e12921.

[36] Barthelemy NR, Li Y, Joseph-Mathurin N, Gordon BA, Hassenstab J, Benzinger TLS, et al. A soluble phosphorylated tau signature links tau, amyloid and the evolution of stages of dominantly inherited Alzheimer’s disease. Nat Med. 2020;26:398–407.

[37] Greenberg SG, Davies P. A preparation of Alzheimer paired helical filaments that displays distinct tau proteins by polyacrylamide gel electrophoresis. Proc Natl Acad Sci U S A. 1990;87:5827–31.

38. Guide for the Care and Use of Laboratory Animals. 8th ed. Washington (DC)2011.

[39] He Z, McBride JD, Xu H, Changolkar L, Kim SJ, Zhang B, et al. Transmission of tauopathy strains is independent of their isoform composition. Nat Commun. 2020;11:7.

[40] Wirths O. Preparation of Crude Synaptosomal Fractions from Mouse Brains and Spinal Cords. Bio Protoc. 2017;7:e2423.

[41] Seyfried NT, Dammer EB, Swarup V, Nandakumar D, Duong DM, Yin L, et al. A Multi-network Approach Identifies Protein-Specific Co-expression in Asymptomatic and Symptomatic Alzheimer’s Disease. Cell Syst. 2017;4:60–72 e4.

[42] Ping L, Duong DM, Yin L, Gearing M, Lah JJ, Levey AI, et al. Global quantitative analysis of the human brain proteome in Alzheimer’s and Parkinson’s Disease. Sci Data. 2018;5:180036.

[43] Zhou M, Duong DM, Johnson ECB, Dai J, Lah JJ, Levey AI, et al. Mass Spectrometry-Based Quantification of Tau in Human Cerebrospinal Fluid Using a Complementary Tryptic Peptide Standard. J Proteome Res. 2019;18:2422–32.

[44] Zhou M, Haque RU, Dammer EB, Duong DM, Ping L, Johnson ECB, et al. Targeted mass spectrometry to quantify brain-derived cerebrospinal fluid biomarkers in Alzheimer’s disease. Clin Proteomics. 2020;17:19.

[45] Brademan DR, Riley NM, Kwiecien NW, Coon JJ. Interactive Peptide Spectral Annotator: A Versatile Web-based Tool for Proteomic Applications. Mol Cell Proteomics. 2019;18:S193–S201.

[46] Iqbal K, Gong CX, Liu F. Hyperphosphorylation-induced tau oligomers. Front Neurol. 2013;4:112.

[47] Iqbal K, Liu F, Gong CX, Alonso Adel C, Grundke-Iqbal I. Mechanisms of tau-induced neurodegeneration. Acta Neuropathol. 2009;118:53–69.

[48] Iqbal K, Grundke-Iqbal I. Discoveries of tau, abnormally hyperphosphorylated tau and others of neurofibrillary degeneration: a personal historical perspective. J Alzheimers Dis. 2006;9:219–42.

[49] Wood JG, Mirra SS, Pollock NJ, Binder LI. Neurofibrillary tangles of Alzheimer disease share antigenic determinants with the axonal microtubule-associated protein tau (tau). Proc Natl Acad Sci U S A. 1986;83:4040–3.

[50] Grundke-Iqbal I, Iqbal K, Quinlan M, Tung YC, Zaidi MS, Wisniewski HM. Microtubule-associated protein tau. A component of Alzheimer paired helical filaments. J Biol Chem. 1986;261:6084–9.

[51] Delacourte A, Defossez A. Alzheimer’s disease: Tau proteins, the promoting factors of microtubule assembly, are major components of paired helical filaments. J Neurol Sci. 1986;76:173–86.

[52] Vogel JW, Iturria-Medina Y, Strandberg OT, Smith R, Levitis E, Evans AC, et al. Spread of pathological tau proteins through communicating neurons in human Alzheimer’s disease. Nat Commun. 2020;11:2612.

[53] Oikawa N, Kimura N, Yanagisawa K. Alzheimer-type tau pathology in advanced aged nonhuman primate brains harboring substantial amyloid deposition. Brain Res. 2010;1315:137–49.

[54] Cramer PE, Gentzel RC, Tanis KQ, Vardigan J, Wang Y, Connolly B, et al. Aging African green monkeys manifest transcriptional, pathological, and cognitive hallmarks of human Alzheimer’s disease. Neurobiol Aging. 2018;64:92–106.

[55] Schultz C, Hubbard GB, Rub U, Braak E, Braak H. Age-related progression of tau pathology in brains of baboons. Neurobiol Aging. 2000;21:905–12.

[56] Edler MK, Munger EL, Meindl RS, Hopkins WD, Ely JJ, Erwin JM, et al. Neuron loss associated with age but not Alzheimer’s disease pathology in the chimpanzee brain. Philos Trans R Soc Lond B Biol Sci. 2020;375:20190619.

[57] Alonso A, Zaidi T, Novak M, Grundke-Iqbal I, Iqbal K. Hyperphosphorylation induces self-assembly of tau into tangles of paired helical filaments/straight filaments. Proc Natl Acad Sci U S A. 2001;98:6923–8.

[58] Goedert M, Spillantini MG, Cairns NJ, Crowther RA. Tau proteins of Alzheimer paired helical filaments: abnormal phosphorylation of all six brain isoforms. Neuron. 1992;8:159–68.

[59] Barthelemy NR, Saef B, Li Y, Gordon BA, He Y, Horie K, et al. CSF tau phosphorylation occupancies at T217 and T205 represent improved biomarkers of amyloid and tau pathology in Alzheimer’s disease. Nat Aging. 2023;3:391–401.

