## Supplementary methods and data for "Marmosets as model systems for the study of Alzheimer’s disease and related dementias: substantiation of physiological Tau 3R and 4R isoform expression and phosphorylation"

### Supplemental Methods

#### 1.1 qPCR quantification of MAPT mRNA levels

Quantitative PCR (qPCR) was conducted using the SYBRgreen-based method by PowerUp™ and SYBR™ Green Master Mix, a qPCR thermocycler (Applied bioscience). Equal amounts of reverse-transcribed cDNA from the brain tissue of marmosets of the following ages were used as a template: one subject at postnatal day 1, one adolescent subject aged 12 months, and one aged adult of 9 years. For the 3R targeted reaction, the following 3R Tau-specific primers were used: Forward primer: GAAGGTGCAAATAGTCTACAA; reverse primer: GAGACCCGATCTTCGACTGGAC. PPIA primer was used as the control to normalize the target mRNA expression levels: forward primer: TGCTGGACCCAACAGAAACGGT; reverse primer: AAAGCGCTCCATGGCCTCCA. Each reaction was carried out in triplicate in separate wells in a total volume of 10 μL, with 1 μL of reverse-transcribed cDNA template (1 ng/μL). A reagent without a cDNA template was included in each PCR reaction to ensure no contamination from the master mix. The relative quantity of each reaction was normalized to the relative amount of PPIA mRNA. For analysis, the mean of each triplicate reaction was averaged as the final quantity of the sample, and the formula for fold-change ($2^{-\Delta\Delta Ct}$) was used to analyze the mRNA expression level of the targeted genes.

#### 1.2 Immunohistochemistry (IHC)

Tissue preparation was described in Materials and Method 2.2. Paraffin-embedded 4μm brain sections were used for IHC and Mouse on Mouse (M.O.M.) Elite Immunodetection Kit (Vector laboratories #PK-2200) was used for immunostaining. The sections were deparaffinized and dehydrated, followed by heat-induced antigen retrieval, then incubated in 0.3% $H_2O_2$ at room temperature for 15 min to block endogenous peroxidase activity. After incubation with 0.1% TritonX100 for 10 min at room temperature, the slices were blocked with blocking buffer: 1% horse serum, 5% goat serum, 5% donkey serum diluted in TBS (150 mM NaCl, 50 mM Tris, pH 7.6) with 0.1% TritonX100) at room temperature for 1hr, followed by incubation with primary antibodies diluted in blocking buffer at 4 °C overnight. After washing by TBST (TBS with 0.1% Tween20), the slices were immersed in 1% horse serum in TBS-T containing biotinylated secondary antibodies (Vector Laboratories, Inc., Burlingame, CA) for 45 min. Bound antibodies were labeled with avidin and biotinylated HRP (VECTASTAIN Elite ABC kit; Vector Laboratories, Inc.) and developed with 3,3-diaminobenzidine (DAB) substrate (Vector Laboratories #SK-4100). Slices were mounted with VectaMount Permanent Mounting Medium (Vector laboratories#H-5000) and imaged by Zeiss Axiovert 200M microscope. Mouse anti 3Rtau antibody RD3 (1:200) and biotinylated anti-mouse secondary antibody were used for 3Rtau staining. Rabbit anti-4Rtau antibody (1:300) (Cell Signaling#79327) and biotinylated anti-rabbit secondary antibody were used for 4Rtau staining.

**Table 1. Demographics of Marmosets tissues analyzed in the present studies.**

| Subject ID | Sex | Age at Tissue Collection | Analysis and Figure |
|---|---|---|---|
| AP | female | Neonate (Postnatal Day 1) | Western blot (Fig 1B&C, 5)<br>Mass spectrometry (Fig 2,3)<br>RT-PCR (Fig 1A, S2), qPCR (Fig S1) |
| MO | male | 12 months | Western blot (Fig 1B&C, 5)<br>Mass spectrometry (Fig 2, 3) |
| WI | female | 13 months | Western blot (Fig 1B&C, 5A)<br>Mass spectrometry (Fig 2,3) |
| VI | female | 13 months | Western blot (Fig1B&C, 5A)<br>Mass spectrometry (Fig 2, 3)<br>RT-PCR (Fig1 A, S2), qPCR (Fig S1) |
| HU | male | 7 years | Western blot (Fig 1B&C, 4, 5)<br>Mass spectrometry (Fig 2,3) |
| AR | female | 7 years | Western blot (Fig1B&C, 4, 5)<br>Mass spectrometry (Fig 2, 3) |
| ST | female | 9 years | Western blot (Fig 1B&C, 4, 5)<br>Mass spectrometry (Fig Y)<br>RT-PCR (Fig1 A, S2), qPCR (Fig S1) |
| CO | male | 12 years | Immunofluorescence staining (Fig. 1D, Fig. S3)<br>Immunohistochemistry staining (Fig. S4) |

**Table 2.** Post-mortem frozen human brain of Alzheimer's Disease and non-demented control sections of the frontal cortex were obtained from the Emory Alzheimer's Disease Research Center brain bank (pathological traits described in supplemental table #2), used for Mass spectrometry.

| | | Sample size | |
|---|---|---|---|
| | | Control | AD |
| | | *N* = 4 | *N* = 4 |
| **Characteristics** | | | |
| **Age:** | years ± SD | 73 ± 4.24 | 67.5 ± 5.26 |
| **Sex:** | N Males | 2 | 2 |
| | N Females | 2 | 2 |
| **Race:** | N Caucasian | 3 | 4 |
| | N Hispanic | 1 | 0 |
| **PMI:** | hrs ± SD | 7 ± 3.08 | 5.25 ± 1.55 |
| **Amyloid burden:** | CERAD 0-1 | 4 | 0 |
| | CERAD 2-3 | 0 | 4 |
| **Tau burden:** | Braak 0-II | 4 | 0 |
| | Braak III-VI | 0 | 4 |
| ***APOE genotype:*** | *E33* | 4 | 0 |
| | *E34* | 0 | 4 |

**Abbreviations**: AD, Alzheimer's disease
PMI, postmortem interval; *APOE* , apolipoprotein E

### Supplemental Figures

**Fig S1**

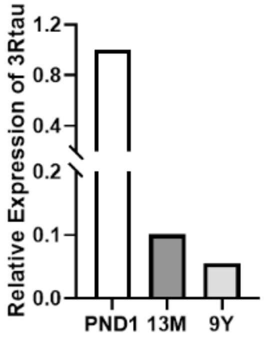


**Fig S1**. **Quantification 3R Tau expression in marmoset brain across the lifespan.** The mRNA expression level without exon 10 (3Rtau) was quantified by qPCR from three different marmosets spanning the following age groups: Postnatal day 1 (PND1), adolescent (13-months; 13M), and aged adult (9 years, 9Y). 3R Tau mRNA expression was 10-15-fold higher in neonate marmoset brains relative to adolescent and aged adult marmoset brain.

**Fig S2**

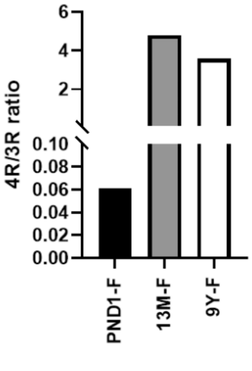


**Fig S2. Ratio of 4R/3R tau mRNA in marmosets of different ages.** The 4Rtau/3Rtau mRNA ratio was calculated from three unrelated marmosets at different ages. The optical density (OD) of each band was quantified using ImageJ software. The OD ratio with exon 10 was calculated relative to the OD ratio without exon 10 ratio. PND1-F, female marmoset at postnatal day 1; 13M-F, female marmoset aged 13 months; 9Y-F, female marmoset aged 9 years.

Fig S3

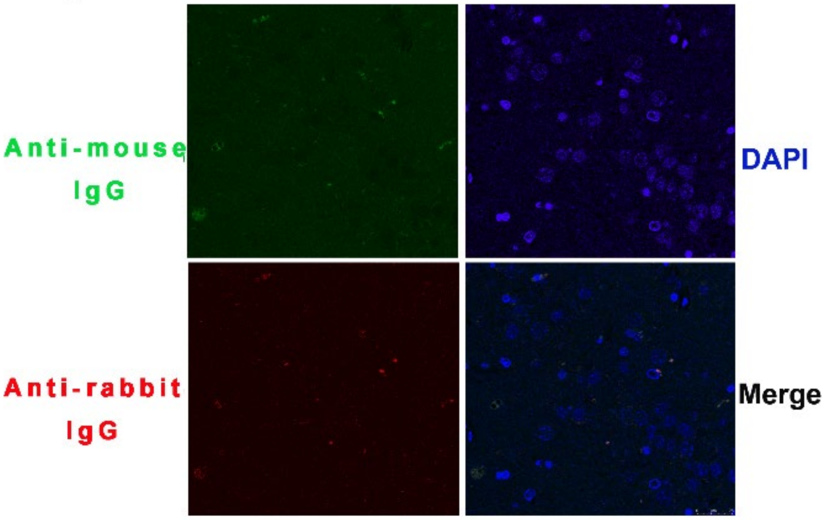


**Fig S3. Negative control staining of IF.** Representative confocal images of background staining, in the absence of incubation with primary antibodies, for the images presented in Figure 1D. A 12-year aged male marmoset brain section was stained with Alexa fluor 488 (Green) conjugated anti-rabbit IgG and Alexa fluor 647 (Red) conjugated anti-mouse IgG, without incubation with the primary antibodies RD3 and anti-4Rtau antibody. Cell nuclei were stained with DAPI (blue). Scale bars are 25 µm.

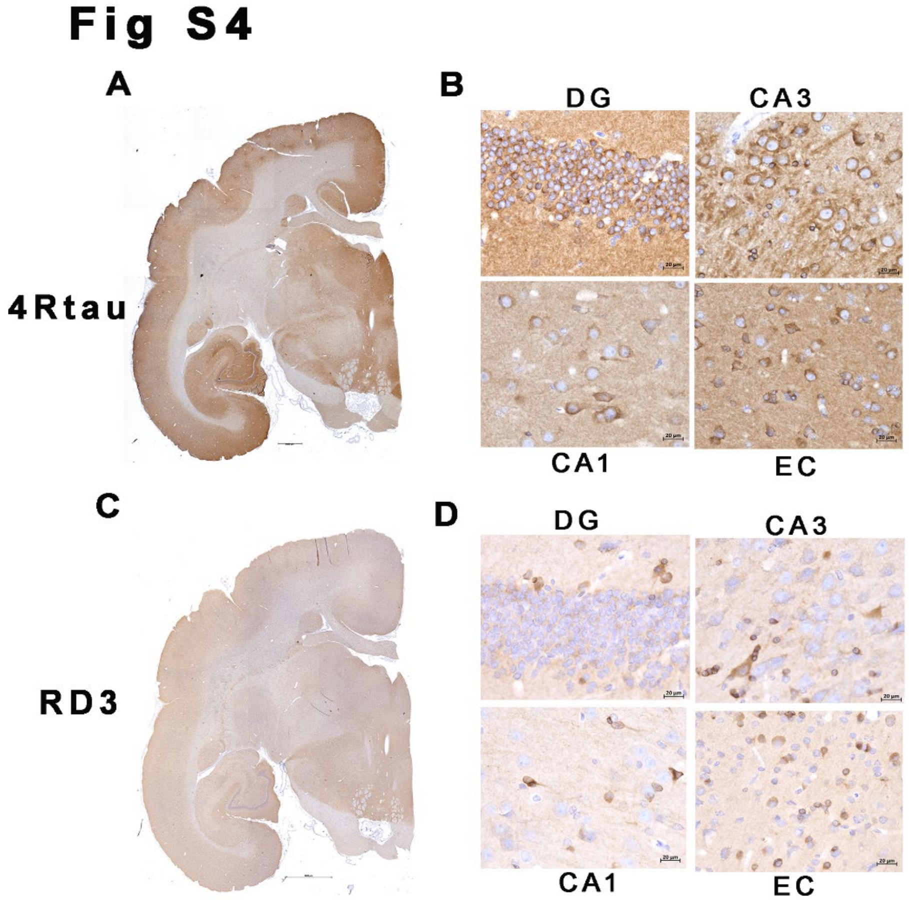


**Fig S4. Immunohistochemistry of 12-year aged marmoset brain.** As a companion to the immunofluorescence staining of the same subject as presented in Figure 1D, paraffin-embedded slices from a 12-year-old male marmoset brain were stained with anti-3RTau (RD3) and anti-4RTau antibodies with HRP conjugated secondary antibody. **A and B)** 4Rtau immunostaining images in marmoset brain. The image was obtained with 2.5x objective lens, scale bar 1000 µm (**A**)**.** Representative magnified images from Hip and EC regions were obtained using a 40x objective lens, scale bar 20µm (**B**). **C and D)** 3Rtau expression detected by 3Rtau specific antibody RD3 in marmoset brain. The image was obtained with 2.5x objective lens, scale bar 1000µm(C)**.** Representative magnified images from Hip and EC regions were obtained by a 40x objective lens, scale bar 20µm (D). Cell nuclei were stained with hematoxylin (blue). CA1 is hippocampus CA1 area, CA3 is hippocampus CA3 region, DG is the hippocampus dentate gyrus region, and EC is the entorhinal cortex area.
